# WormAI: Artificial Intelligence Networks for Nematode Phenotyping

**DOI:** 10.1101/2025.02.06.636827

**Authors:** Maggie Lieu, Qi Cao, Ruoqing Meng, Dong Zhang, Veeren M. Chauhan

## Abstract

The nematode *Caenorhabditis elegans* is a key model organism in biological research due to its genetic similarity to humans and its utility in studying complex processes. Traditional image analysis methods, such as those using ImageJ, are labour-intensive, which has led to the integration of AI. This study introduces an AI framework with three machine learning models: WormGAN, a Generative Adversarial Network for generating synthetic nematode images to enhance training data; WormDET, for precise movement tracking; and WormREG, for accurate anatomical measurements. Together, these tools significantly improve the efficiency and accuracy of phenotypic analysis. WormAI demonstrates substantial potential for high-throughput dataset analysis, advancing research in systems biology, drug discovery, and aging. This framework streamlines workflows, enabling faster and more precise discoveries in *C. elegans* studies.

---

*Caenorhabditis elegans* has long served as a cornerstone in biological research, owing to its genetic and molecular similarity to humans [2]. Despite their evolutionary distance, the high degree of gene homology between *C. elegans* and humans [37] has enabled researchers to explore the cellular and molecular mechanisms underpinning complex biological processes, such as systems biology [34] and organismal development [15]. Due to its simple, transparent body structure, well-characterised genetic makeup, and ease of cultivation it has proven to be invaluable in fields like neurobiology and genetics [9]. The organism’s rapid life cycle, spanning approximately three days from egg to adult [3], permits the observation of multiple generations within short timeframes, thereby accelerating research cycles. Furthermore, the transparency of the nematode [5] facilitates the direct observation of internal structures, aiding the study of physiological changes [6].

However, the study of *C. elegans* physiology and behaviour, particularly across diverse environments using conventional microscopy data, presents challenges [21, 18, 33]. While several software tools exist for quantifying *C. elegans* phenotypes from still images, such as `WormSizer` [24], `QuantWorm` [17], and `WormToolbox` [36], these programs often require substantial manual input and are typically designed for specific research needs. Traditional image analysis tools, such as manual observation and the use of open-source software like `ImageJ`^1^ or `Fiji`, can be labour-intensive, time-consuming, and lack robustness in handling large datasets [13, 28]. Therefore, the integration of artificial intelligence (AI) has emerged as a promising solution to these challenges [27, 12, 1]. The OpenWorm project [32], for instance, exemplifies AI’s potential by digitally reconstructing *C. elegans* to simulate its neural and muscular systems at a cellular level, paving the way for further AI-driven research into the worm’s neural circuitry and physical behaviour.

Within the AI domain, machine learning, particularly Convolutional Neural Networks (CNNs) [19, 23] and Generative Adversarial Networks (GANs) [11, 7], offer powerful ways for the rapid and consistent analysis of large volumes of visual data. These technologies have proven especially useful in three key areas: 1) accurately tracking and quantifying movement patterns [40, 22, 30], 2) automatically analysing images and videos to measure growth parameters (phenotypic quantification) [12, 31, 4], and 3) identifying patterns in extensive datasets to formulate new hypotheses about biological mechanisms. CNNs excel in image recognition and classification tasks [38], making them ideal for identifying and classifying the various life stages of *C. elegans*, analysing morphological changes, and detecting subtle behavioural variations. Therefore, the automation of image analysis, CNNs can significantly enhance the efficiency and accuracy of phenotypic quantification.

GANs, on the other hand, are valuable for generating realistic synthetic images of objects of interest, such as nematodes [14, 39]. This capability is particularly important for augmenting limited training data, improving the robustness of machine learning models, and enabling the simulation of hypothetical scenarios to explore the effects of different environmental conditions on the nematodes *in silico*. Therefore, integrating GANs and CNNs into a unified analysis pipeline potentially provides a comprehensive approach to visual data processing, enabling detailed studies of *C. elegans* at a scale and precision that are unattainable through manual methods. Despite these advancements, the field still faces challenges in ensuring data quality, developing algorithms capable of generalising across diverse datasets, and managing computational demands [1]. Addressing these challenges involves rigorous data preprocessing, the development of specialised neural network architectures, and the use of high-performance computing resources.

In this research, we developed a set of novel machine learning methods for the rapid analysis of large datasets of *C. elegans* images. Our approach, once trained, will accurately recognise and localise nematodes, allowing for precise measurement of anatomical dimensions such as length, width, and volume at each life cycle stage. Furthermore, this method will facilitate the tracking of nematode movement between frames, supporting comprehensive behavioural and phenotypic analysis. By creating *in silico* models that simulate life and whole organism biology, this approach holds the potential to provide deeper insights into biological mechanisms and foster new discoveries.

## Results

To generate and analyse synthetic nematodes, we developed a multi-stage computational pipeline comprising three key components: WormGAN, WormDET, and WormREG. (Figure 1). This framework enables the realistic simulation, detection, and regression of nematode characteristics, leveraging deep learning techniques to automate and enhance nematode analysis.

**Fig. 1.**
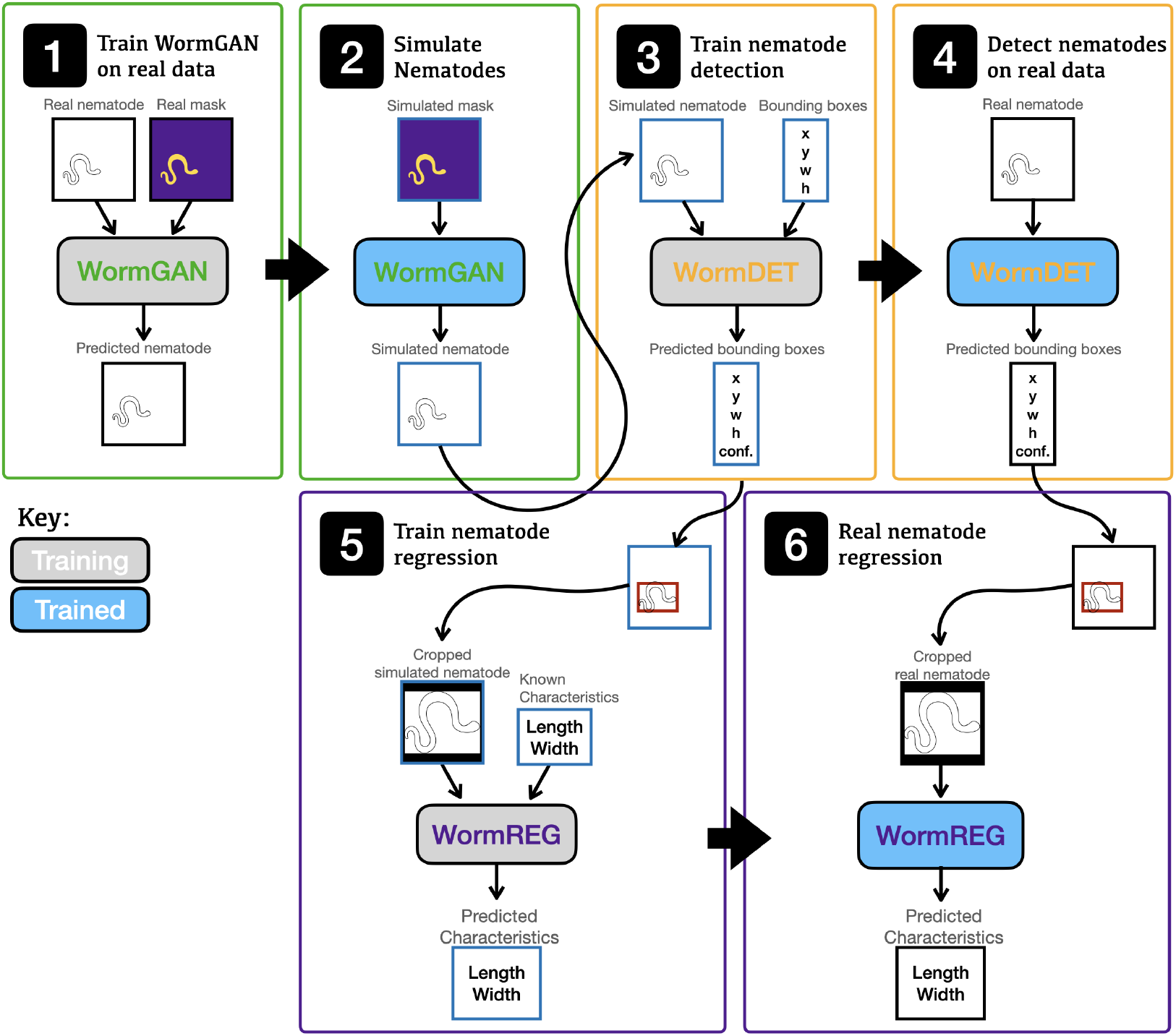
WormAI pipeline workflow. The 3 models, WormGAN, WormDET and WormREG are color coded green, orange and purple respectively. The training stage of the models where the parameters are being updated is shaded grey and the trained models, where the parameters are fixed are shaded blue.

This pipeline facilitates the creation of a large-scale dataset of synthetic nematodes, which is then used to train and validate detection and regression models. This permitted the generation of realistic nematode images, accurately detecting nematodes, and predicting their morphological properties.

### Synthetic Nematode Generation

Using WormGAN, we successfully generated a dataset of 10,000 synthetic nematode images (Figure 2). Each image was conditioned on semantic segmentation maps created using our defined parametric approach, ensuring that the generated nematodes maintained realistic length and width distributions. The GAN model, trained on 94 real image-segmentation pairs, was able to generate highly detailed and visually coherent nematode structures.

**Fig. 2.**
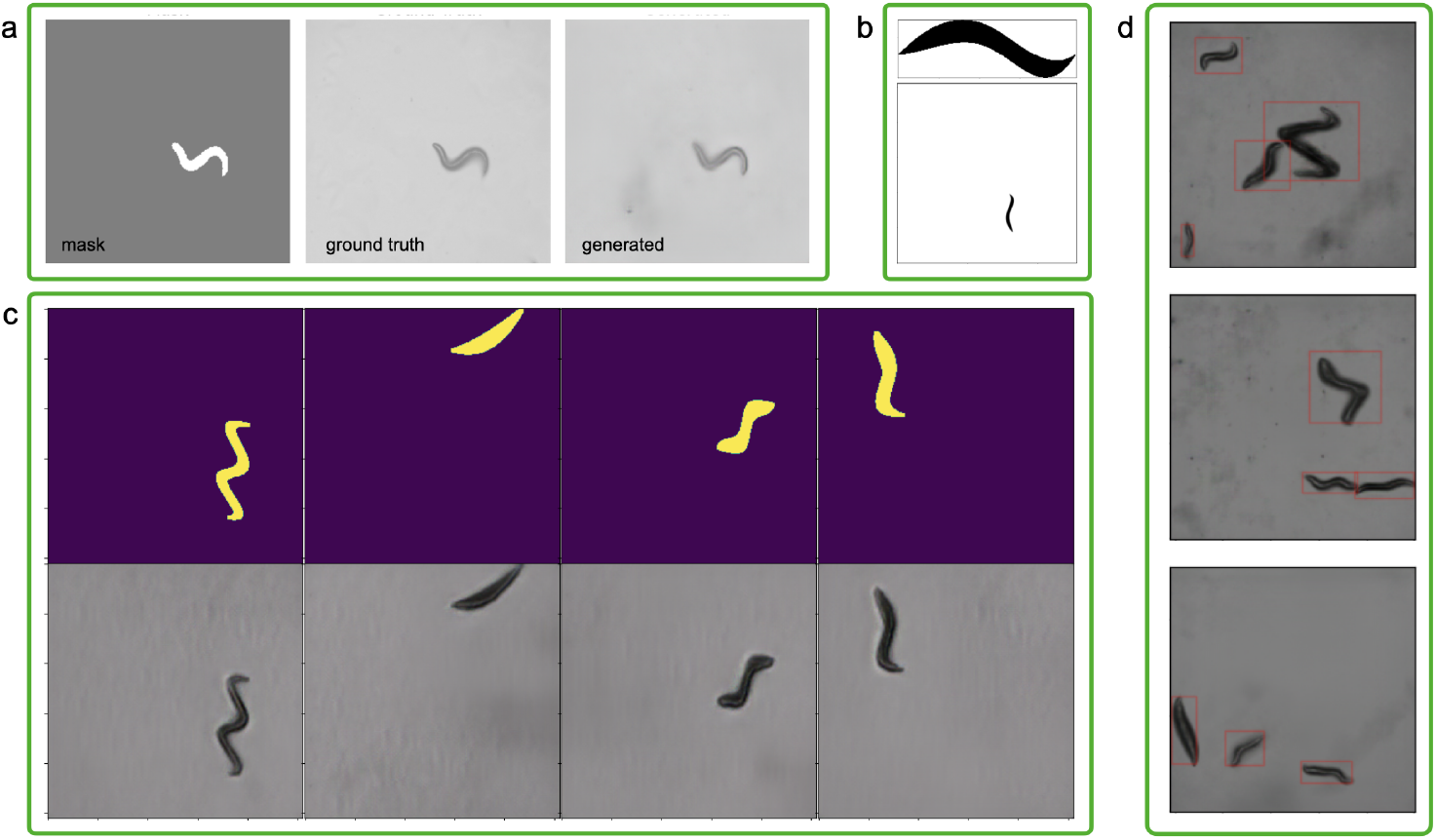
WormGAN synthsising simulated nematodes for analysis. *a:* Epoch 15 of GAN training. From left to right is the input mask, the ground truth label image and the GAN generated image. *b:* Cropped segmentation map of simulated worm and full segmentation map of simulated nematode. *c:* GAN simulated nematodes with the corresponding input masks. *d:* Simulations of multiple nematodes with 4, 3, and 3 nematodes localised within their known bounding boxes (red).

To assess the realism of the generated nematodes, we visually inspected a subset of images and compared their morphological features to real nematodes. The synthetic worms closely matched the expected shape, curvature, and segmentation features, demonstrating that the model successfully preserved key characteristics from the training data. Additionally, the synthetic nematode dataset displayed a range of natural variations in undulation patterns and body widths, which were induced by the procedural segmentation map generation process.

To ensure that the synthetic dataset accurately represented real nematodes, we compared the length and width distributions of the generated nematodes to the original dataset. The linear regression model used to determine nematode width as a function of length yielded results that closely matched the distribution of real nematodes, with minimal deviation. The addition of normally distributed noise (*σ* = 8.36) helped account for natural variations observed in real data. The Student-t distributed displacement function for undulations also introduced biologically plausible variations in nematode shape.

To create a dataset that better resembled real-world microscopy images, we generated composite images containing multiple simulated nematodes (Figure 2d). Each image contained between 0 and 6 nematodes, selected randomly from the single-worm dataset. This approach resulted in a dataset of 10,000 composite images, effectively simulating the clustering and density variations seen in real samples. The final dataset thus provides a diverse and scalable alternative to manually captured images, significantly reducing the time and effort required for nematode imaging and measurement.

Overall, our method demonstrates the effectiveness of GAN-based synthetic nematode generation, providing a scalable, high-fidelity alternative to manual imaging while maintaining morphological accuracy and variability.

### Nematode detection

The nematode detection model trained in ~6 hours on 2 NVIDIA RTX A6000s. On completion of training, the inference time was ~ 0.1 seconds for a single image.

Figure 3 shows a examples of the test data with bounding box and confidence score overlays from the model and the ground truth demonstrating agreement. Our network achieved a mean Intersection over Union (IoU) score of 87.6% on the test dataset. This was consistent visually, such that the predicted boxes match the ground truth boxes with high confidence. It was also confirmed that the network was particularly competent at assigned the correct number of boxes to the number of nematodes present in each image. This was also true for boxes that did not demonstrate overlapping. This tends to be the case where there are multiple worms with significant overlap.

**Fig. 3.**
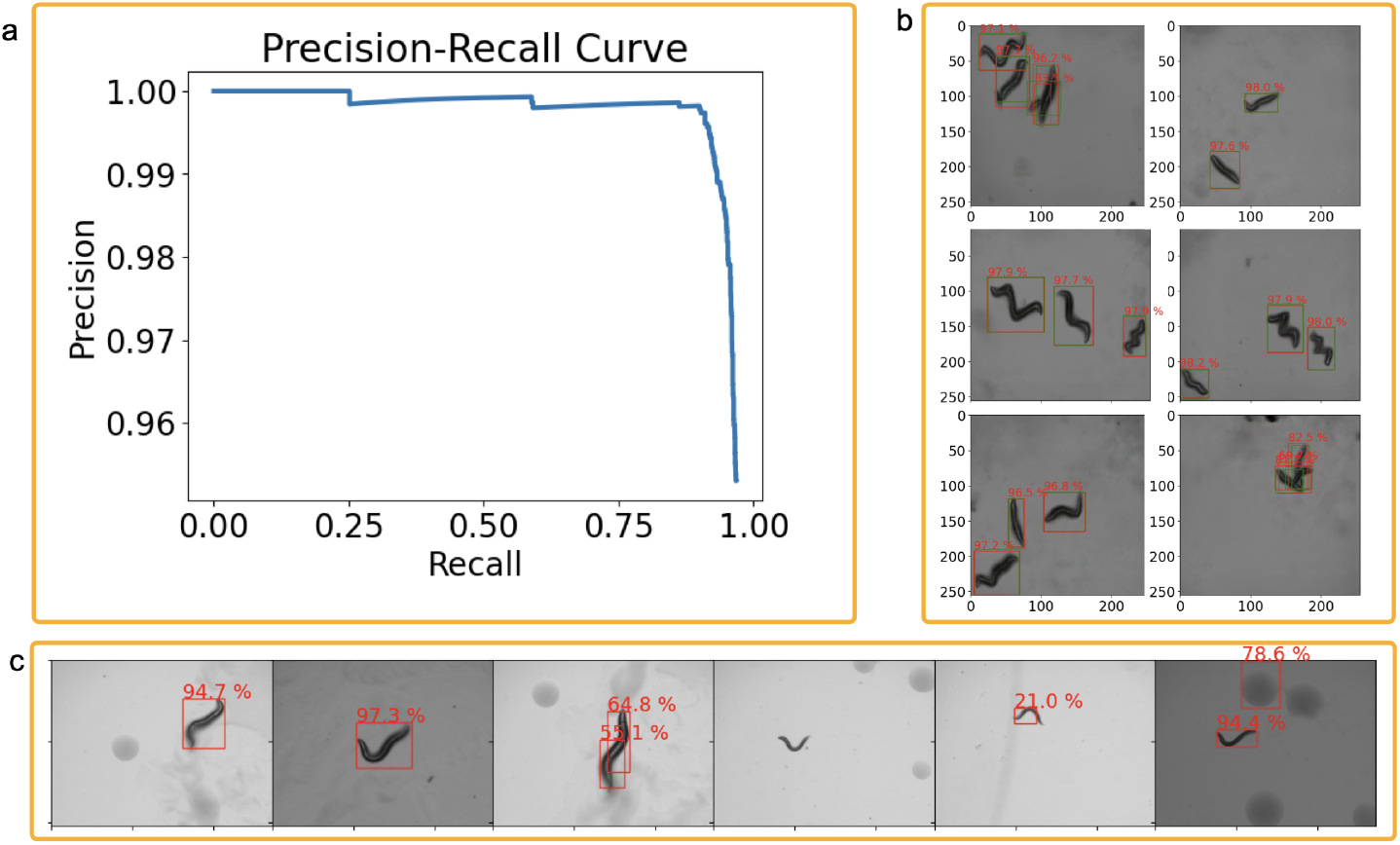
Nematode detection using WormDET. *a:* Precision Recall curve of the WormDET model at an IoU threshold of 0.5. *b:* Localizations on the example test dataset images of simulated nematodes. The green boxes correspond to the ground truth. The red boxes are detections with detection probability from the model. Axes values are in pixels. *c:* Examples of the detection model on real data showing a mixture of true positives, false positives, false negatives and low confidence scores.

The network achieves precision and recall scores at an IoU threshold of 0.5, of 95.7% and 97.2% respectively, where precision measures how many of the detected objects are correct, while recall measures the number of objects that were correctly detected. These metrics provide insights into the trade-off between false positives and false negatives. Figure 3a is the precision-recall curve that shows the trade-off between precision and recall at different detections confidence thresholds. The Average Precision (AP) summarises the precision-recall curve into a single value, which represents the average of all precisions at different recall levels. The AP is calculated as the area under the precision-recall curve. Our network achieves an AP score of 97.0%, where a higher AP indicates an better-performing model, as it means the model achieves both high precision and high recall across various confidence levels. To put this into perspective, the state-of-the-art model on the Hugging Face object detection leader-board^2^ as of January 2025 achieved a mean average precision of 72.1% over many classes of detections.

WormDET exhibited challenges in detecting shorter and thinner nematodes, which were attributed to biases produced by the WormGAN dataset. We hypothesise that the inclusion of a more diverse sample of training data could further improve WormDET performance.

### Applied to real data

Applied on the real data (Figure 3c) the nematode detection model reconfirms what we found in the simulation test data, notably that smaller nematodes tended to be missed, or were detected with low confidence. A few larger nematodes had duplicate detections. These likely arose from the model identifying multiple features within the larger nematodes, triggering overlapping bounding boxes. To address this, non-maximum suppression (NMS), a common technique in object detection, can be employed [10]. NMS filters multiple bounding boxes that overlap significantly, retaining only the box with the highest confidence score. Typically an IOU score of 50% is used to define duplicates however the duplicate detection in the third panel of Figure 3c has an IOU score of 39% so a smaller threshold would be needed to eliminate the duplicate, in this particular case. This would have a trade off of missing worms that are close in proximity.

### Nematode Regression

The simulated nematodes were used to train a regression model to predict the width and length of nematodes from isolated nematode cutouts, which were produced using the bounding boxes predicted by WormDET. Figure 4a shows the recovery of the parameters in the test set consisting of 2529 simulated worms, where the mean fractional error on length and width are 0.039 and 0.066, respectively. The model consistently assigns accurate length and widths. It is important to note that there were predicted overestimation towards the longer length nematodes. However, it was unclear the significance of this due to the low number of simulated nematode samples in the data range with true length greater than 750 px. It was anticipated that biases would be observed at the lower true widths and lengths, due to the nature of the final activation, however these were not apparent.

**Fig. 4.**
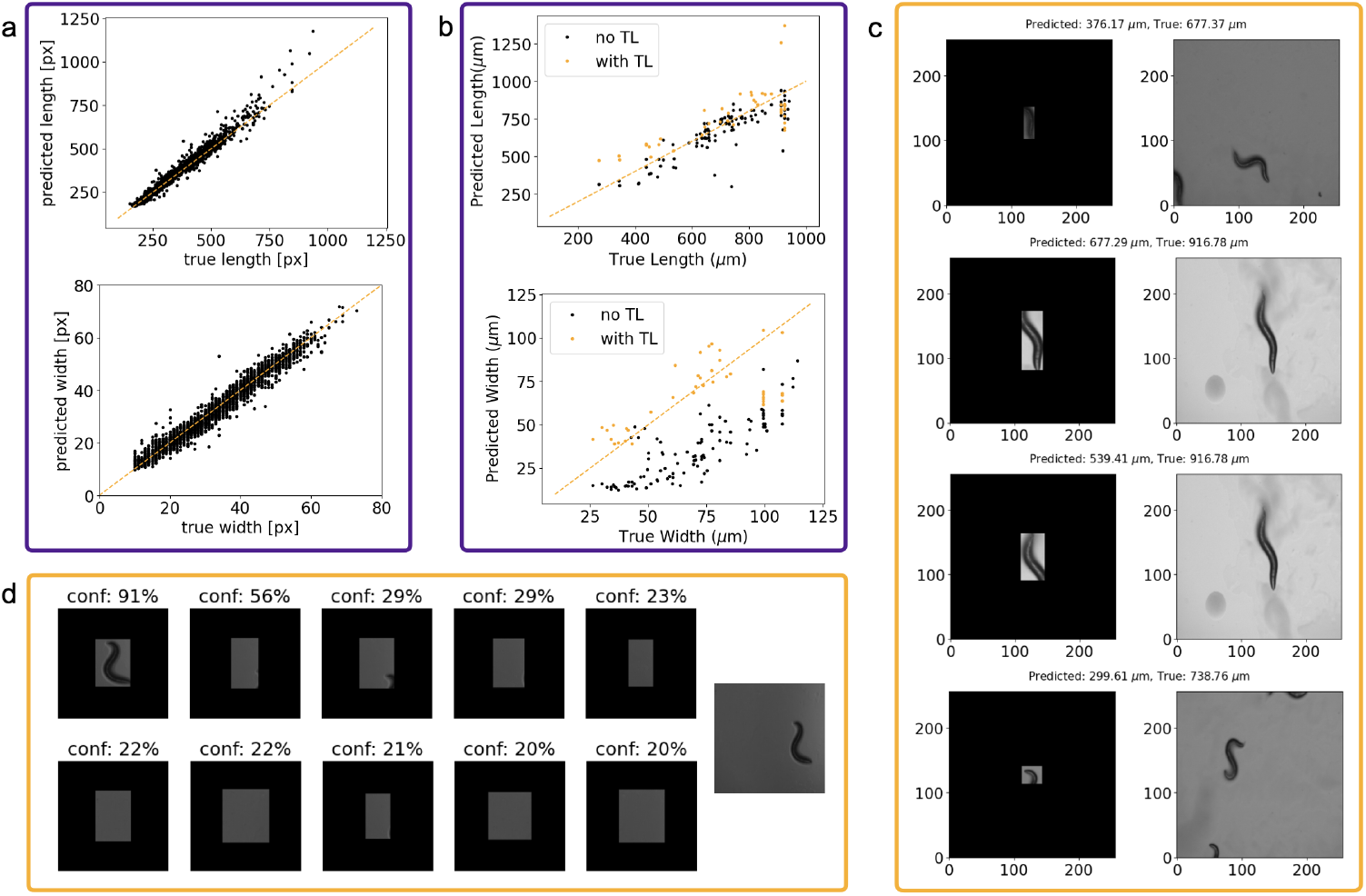
Exploration of WormREG combined with WormDET. *a:* Regression model predictions of width and length of simulated nematodes. *b:* Regression model predictions of width and length of real nematodes before and after fine tuning the model with transfer learning [25]. *c:* Outliers in the length prediction of real nematodes. Left images are the extracted cutouts from the object detection model, right images are the corresponding full image. These can be attributed to multiple detections and detections off the edge of the image. The incorrect box predictions lead to underestimated length predictions as listed in the main text of the images. The images are 256 *×* 256 pixels. *d:* False positive detections from the object detection model, play a significant role in the accuracy of the width and length prediction. This shows the 10 detections made on the input image shown on the right only has 1 worm. The confidence of the detections is shown above.

We used this regression model on all positive detections in the real data sample, creating cutouts from the bounding boxes to predict the nematode characteristics. The mean fractional error on the on length and width were 0.110 and 1.170, respectively. The length predictions are consistent with a few notable outliers. On visual inspection, its becomes apparent that these are due to either duplicate detections of a single nematode, or nematodes that have been detected on the edge of the image and are not the subject of interest (Figure 4c). The width predictions exhibited a underestimation of true values and further calibration was necessary to achieve comparable accuracy for width. This was achieved, through the application of transfer learning, fine-tuning the model with an additional 25 epochs of training on 50% of the real nematode data, which was composed of 94 detections with a reservation of 10% of data for validation. Transfer learning improved mean fractional error for width by 450% to 0.259, on the held-out test set, with marginal influence to length (0.149). This significantly mitigated the underestimations previously observed in width predictions. A subtle reduction in length prediction accuracy was identified, however this evaluation was conducted on only 38 nematodes, compared to 94 without transfer learning, and therefore was not directly comparable to the initial results.

There were also some outliers, in particular Figure 4b highlights sets of vertical points with the same true length values, e.g., at 916 *μ*m, which were multiple detections on the same image (Figure 4c). Figure 4d shows such an example where many false positive detections are made on the same image from the object detection model. However, the confidence of the false positives were low, and choosing a suitable detection threshold would remove many of these. At a threshold of 50% confidence, the fractional error on length and width reduces to 0.140 and 0.143 respectively. Adjusting the confidence score threshold allows users to fine-tune the nematode detection algorithm to better suit their needs. Lowering the threshold increases the chance of detecting all nematodes (higher completeness), but may also lead to the inclusion of false positives (lower purity). Raising the threshold ensures that only high-confidence detections are reported (higher purity), but some true nematodes might be missed (lower completeness). Users can choose a threshold that balances these factors based on the specific requirements of their task.

## Discussion

In this study we present the WormAI framework and show that it demonstrates significant advancements in automating and improving the phenotypic analysis of *C. elegans* through a generative adversarial network (WormGAN), a detection network (WormDET), and an inference network (WormREG). WormGAN successfully generates realistic synthetic images of *C. elegans*, to augment the training datasets that are otherwise limited by the labour-intensive nature of manual data collection. WormGAN-generated nematodes closely mimic the morphological variability found in real populations. These synthetic nematodes enhance the robustness of models trained for tasks such as detection and regression, reducing biases and errors associated with small or imbalanced datasets.

WormDET achieves a mean Intersection over Union (IoU) score of 87.6% and a high average precision (AP) score of 97.0%, outperforming global benchmarks in similar applications. Despite these strong metrics, WormDET exhibits a notable weakness in detecting the shortest, thinnest worms, leading to a high rate of false negatives. This suggests a potential bias in the WormGAN simulated nematode data or features used for detection. This performance gap does not appear to be influenced by their undulations, indicating that the model effectively captures this morphological characteristic. This is evident from the histograms comparing the distributions of false negatives and true positives (Figure S3) and this highlights a crucial area for improvement in the model’s training and feature engineering, potentially we could simulate more worms with those characteristics to improve this. WormREG accurately predicts nematode length and width, achieving low mean fractional errors for length and width in the simulated test set. When WormReg was applied to real data, additional fine-tuning reduced the errors, highlighting the value of transfer learning in bridging the gap between synthetic and real-world datasets.

WormAI’s ability to simulate *in silico* conditions allows researchers to explore hypothetical scenarios, potentially accelerating studies in systems biology, drug discovery, and aging research. By automating length and width measurements, WormAI reduces the time and resources required for high-throughput phenotypic analysis, paving the way for more efficient experimentation. While WormAI’s performance is impressive, several areas warrant further investigation. Firstly, the detection model’s difficulty with smaller nematodes underscores the need for more representative training datasets, with the synthetic nematodes it would be trivial to adapt this in the future to target these simulations or augmentation strategies to address these gaps. Secondly, though transfer learning improved the model’s accuracy on real data, some biases and outliers remain, particularly in the detection of overlapping or nematodes that encroach on fields of view for analysis. Exploring advanced techniques such as ensemble models or semi-supervised learning might help refine real-world applicability. Lastly, incorporating the WormAI framework into established experimental workflows could provide insights into its practical utility and scalability.

WormAI represents a significant step forward in the integration of machine learning into biological research. We anticipate that its innovation to generate synthetic training data and its automated detection and regression models enhance the precision and efficiency of *C. elegans* phenotyping.

## Methods

### Data

#### Real data collection

*C. elegans* (Bristol N2) were maintained on nematode growth medium (NGM) agar and *Escherichia coli* (OP50) at 20 °C. Mixed populations of nematodes, consisting of larva (L1 - L4) and adults were harvested from culture plates and imaged at 4x magnification using an EVOS microscope equipped with both bright field and differential interference contrast (DIC), captured in an red green blue (RGB) format.

The data set for the analysis conducted in this study consisted of 94 images, 1024 × 1360 px resolution in .tif format, taken at 4x magnification. The image scale is 1.6 *μ*m / pixel and we worked with pixel data unless stated otherwise as is standard in machine learning. Images for analysis contained 1 or more nematodes, and the same nematode may appear in multiple images taken at a different timestamps.

#### Real Labels

We take the traditional approach to measure the lengths and widths of each nematode manually in each image (Figure S1), using the open-source image processing package `Fiji`^3^. This package is based on `ImageJ`^4^ (version 1.53) and bundles together other plugins to facilitate scientific image analysis.

- Nematode length was measured using the segmented line tool in `Fiji` to make a skeletal line along the body of the nematode. The measured value is taken as the sum over the length of each line segment in pixels. The number of segments can vary from nematode to nematode, which can introduce some user based measurement anomalies and whilst this does not give an exact length of the nematode, it provides an appropriate approximation.
- Nematode width was measured using the straight line tool in `Fiji`. Straight lines are made across 5 points along the width of the nematode. The length of these lines in pixels are then averaged.

The measured values are taken as the ground true lengths and widths throughout the rest of this paper. They span 200-1000 *μ*m and 10-120 *μ*m, respectively.

#### Segmentation maps

We manually produced segmentation maps corresponding to each ground truth image (Figure S1c). This trains a network to simulate additional nematodes and later facilitated their localisation. In addition, we increase the image dataset 8-fold by incorporating 90° rotations and vertical and horizontal flips.

### Synthetic nematodes

#### WormGAN

To increase the dataset, we use a Generative Adversarial Network [GAN, 11] trained on image and segmentation map pairs. GANs use two neural networks to compete against each other. One network, called the generator, tries to create new data that is indistinguishable from the real data. The other network, called the discriminator, tries to determine whether the data it sees is real or fake. The network is optimised to minimise a combined loss of both the generator and the discriminator. This competition drives both networks to improve, with the generator getting better at creating realistic data and the discriminator getting better at spotting fakes. Once trained the generator can be used to create highly realistic synthetic data. Here we use GauGAN[26], a GAN model developed by NVIDIA that introduces a new normalization technique called Spatially-Adaptive Normalization (SPADE). This helps preserve semantic information (the labels in the segmentation map) in the generated image, leading to more photo-realistic and coherent results from labeled semantic segmentation maps. Using the standard 80-10-10 train-validation-test split of 94 image-segmentation map pairs, we condition the model on the segmentation maps. The images and segmentation maps are rescaled to size 256 × 256 pixels, a common practice in machine learning to reduce computational cost and improve training stability to aid with convergence. We also used random horizontal and vertical flips to artificially increase the training sample size. The model is trained with a batch size 1 over 25 epochs.

#### Worm simulations

The trained WormGAN requires semantic segmentation maps to produce simulated nematode images. To make the 1000 simulations we create first create the segmentation maps by following the below procedure:

- Sample a random guide length of nematode (*l*^*′*^) and nematode width (*r*). The guide length is an approximation to the real length of the simulated nematode. To maintain a realistic length and width distribution, we fit a linear regression model to the real nematode data. The guide lengths are drawn from a random uniform integer distribution between 150 and 300 pixels,

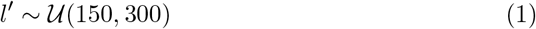

and the widths are computed according to the scaling relation,

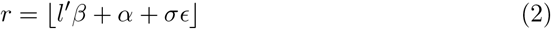

where *ϵ* ~ 𝒩 (0, 1) is a standard normally distributed random variable, and ⌊•⌋ denotes the floor function. The slope and intercept were fit to values *β*=0.12 and *α*=-9.43, respectively. *σ* = 8.36 accounts for the scatter in relation, computed as the standard deviation of the root mean square error (Figure S2).
- Draw a random integer value for the number of segments (*n*_*s*_) that determines the frequency of sinusoidal movement in nematodes.

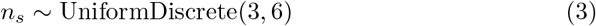

The greater number of segments a nematode has, the larger the freedom of movement.

- To create nematode undulations, the guide length of worm was split into *n*_*s*_ number of segments and at the centre, which were displaced by

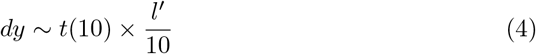

where *t*(*x*) is the student-t distribution with *x* degrees of freedom, which occasionally permits some very large displacements, and the latter component imposes scaling with the length of the worm, as it would be unrealistic for short worms to have very large undulations.
- Interpolate a smooth line over the segments whilst accounting for the worm width using a cubic spline.
- Compute the true length *l* as the euclidean distance between interpolated points (**x, y**).

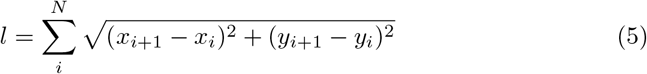
- Randomly rotate the nematode and place at a random location within the 1024×1360 px image corresponding to the size of the real data.

#### Applied WormGAN

The segmentation maps generated are then resized to feed into the trained GauGAN model to produce realistic nematode images with known widths and lengths. This removes the time intensive necessity of manual imaging and measuring their widths and lengths, and also ensures that the characteristics are accurate.

In total this gives us 10,000 simulated nematode images. However we note that each image only contains 1 nematode. In order to create more realistic representation of the ground truth data we create a data sample of multiple simulated worm images from random combinations of the single nematode simulations. To do this, we randomly draw as integer between 0 and 6 to define the number of nematodes per image, and stack that number of randomly selected simulated images to create 10,000 simulated images of multiple nematodes.

#### WormDET

The nematodes are free to roam across the field of view. In order to take accurate size measurements we require centralised images of the nematode of interest. To obtain this, we invoke an object detection model to localise the boxes that encompasses each nematode. The parameters of this are the *x* and *y* coordinates of the box and its height *H* and width *W*. These values are known from the simulations. We explored the use of an object detection model consisting of a simple convolutional neural network [CNN, 20]. Which performed exceptionally on the single nematode data but did not obtain the same results on the simulations where more than 1 nematode is present. Ultimately, we opted for the YOLOv8 detection architecture [35], an improved implementation of YOLO [You Only Look Once 29] developed for high speed object detection and image segmentation. It is a single shot detector where unlike older models that first generate region proposals then classifies them, YOLO performs detection on a single pass of the network employing a convolutional neural network (CNN) to analyse an input image and split the feature maps into grids to predict bounding boxes and class probabilities on objects centered in each grid cell. YOLOv8 builds on YOLO, incorporating a new backbone CNN and removes the use of anchor boxes (predefined box shapes) in favour of keypoint prediction (center prediction and edges) which reduces the number of predictions and thus improves efficiency. Here we employ the smaller network YOLOv8s (13M parameters) and update the weights in batch sizes of 1 images, using an Adam optimiser with a 0.001 learning rate with an exponential decay factor 0.1 on plateau during the training phase. To compare the ground truth bounding boxes to the network predictions, we use the CIoU loss (Complete IoU loss) function. This is an improvement on earlier IoU (intersection over union) metric which measures the amount of overlap between the predicted bounding boxes (*A*) and the ground truth bounding boxes (*B*),

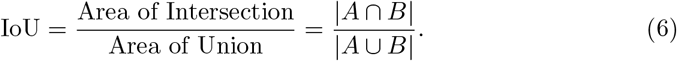

Where boxes don’t overlap, the IoU is 0 and does not provide information on how to move the boxes closer together. It also doesn’t take into account the box centers or their aspect ratios. CIoU loss takes into account a loss for the central separation

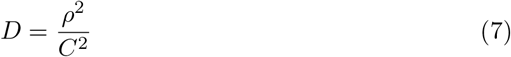

where *ρ* is the Euclidean distance between the center points, and *C* is the diagonal length of the smallest enclosing box covering both *A* and *B*. It also takes into account a loss for the aspect ratio,

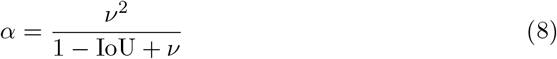

where,

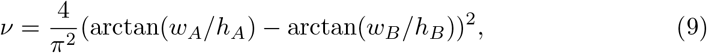

where *w* and *h* correspond to the width and heights of the boxes respectively. The CIoU loss is defined as,

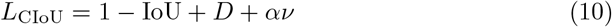

The classification loss for the class of detected object is binary cross entropy,

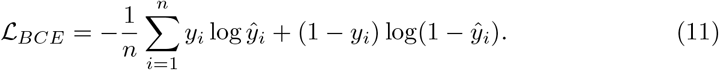

The network was trained on the training data (8000 images) over 50 epochs with early stopping with a patience of 5 to prevent overfitting - where the network was unable to generalise to new data due to it memorising the training data. Early stopping monitors the loss on the validation data (1000 images) to prevent this.

#### WormREG

From the simulations we knew the true length and true width of the simulated nematodes, so, in order to model the lengths and widths of nematodes, we used these as labels to train a CNN to predict them from image crops of the nematodes identified from the nematode detection model.

CNNs use a series of convolution, pooling and dense layers to hierarchically aggregate features in images to (typically) predict properties of the input, in our case, this was the length and widths of the nematodes. Our CNN uses DenseNet121 [16] pretrained on imagenet[8] as the base network. This means the network was first trained on a large number of images in the imagenet dataset to learn to identify patterns within it. The network then leveraged this knowledge, fine-tuning it to a new dataset and problem. This means that we did not need to train the network from scratch saving significant time and resources.

Densenet is an off the shelf architecture that uses dense connections i.e., each layer is connected to every other layer in a feed-forward fashion. Due to this dense connectivity, DenseNet requires fewer parameters than traditional CNNs to achieve similar levels of performance. This is because it can reuse features from all previous layers, reducing the need for learning redundant feature maps. Densenet121 refers to the 121-layer densenet, the smallest of the family just 33MB in size and consisting of 8.1M parameters. Densenet121 obtains a top-5 accuracy score of 92.3% on imagenet classification and therefore we opted for this model over the state-of-the-art densenet201 architecture, which was more than double the size but only improves 1% on the top-5 accuracy.

The last layer of the Densenet121 was removed and replaced by batch normalisation for regularisation and then split into 2 branches, corresponding to the length and width parameters, each with a 50 unit dense layer followed by a 10% dropout. The final activation functions were sigmoid, as the length and widths are normalised by the minmax of the training sample to help the network reach convergence (Figure S4). The input to the network were image cutouts ([256,256,3]) selected from the bounding box detections of the localisation network, these were rescaled, centered and padded to size. We used transfer learning and first train the model for 25 epochs with the Densenet weights frozen to that trained on the imagenet dataset updating the weights in batch sizes of 6 cutouts, with the Adam optimiser and a learning rate of 0.001. These weights were later unfrozen, and trained for a further 25 epochs with a reduced learning rate of 0.0001, a process known as fine-tuning.

Again, we trained the network with early stopping: a patience of 5 epochs to prevent overfitting, and a training-validation-test split of 80:10:10 on 25,285 cutout images. The training data size further increased artificially using horizontal and vertical flip augmentations. We used the piecewise Huber loss to optimise the model, which is a popular choice in regression tasks.

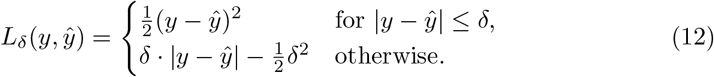

where *δ*=1. It’s evident that the Huber loss is less sensitive to the outliers in the data compared to the squared error loss, as it reverts to the mean absolute error at large error values. Our total loss is the average of the Huber loss from the length and the Huber loss from the width.

Once trained, we further fine-tuned the model by unfreezing the base network weights and continuing training using a learning rate of 0.0001.

## Data availability

Our code and example data are made publicly available at https://github.com/MaggieLieu/wormAI.git

## Acknowledgements

This research utilized computational hardware funded by an STFC Knowledge Exchange Institutional Award (ML). This work was also supported by a Nottingham Research Fellowship from the University of Nottingham (VMC).

## Author Contributions

Conceptualization: M.L., V.M.C., Data Aquisition: V.M.C., Q.C., R. M., D.Z., Software: M.L., Q.C., R.M., D.Z., Formal analysis: M.L., Writing, review and editing: M.L., V.M.C.

## Footnotes

1 https://imagej.net/

2 https://huggingface.co/spaces/hf-vision/object_detection_leaderboard

3 https://fiji.sc/

4 https://imagej.net/

