## Supplementary information for "WormAI: Artificial Intelligence Networks for Nematode Phenotyping"

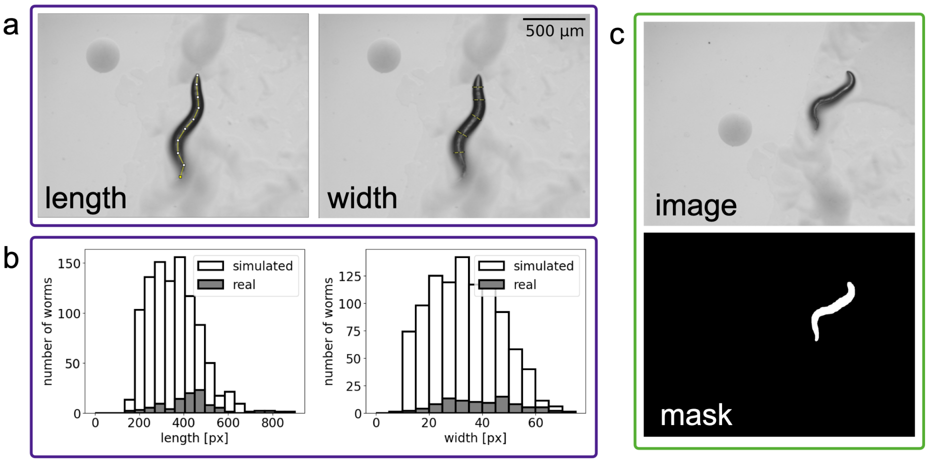


**Fig. S1** *a:* An example image of a nematode with the manually added segmented lines drawn in yellow. Each point represents a a part of a segment which is used to compute the nematode length or to compute the mean nematode width. *b*: The distributions of the measured lengths and widths in the real nematode images and the simulated nematodes. *c:* Input image and its corresponding segmentation map, where pixel values of 1 correspond to nematode pixels and 0 to non-nematode pixels.

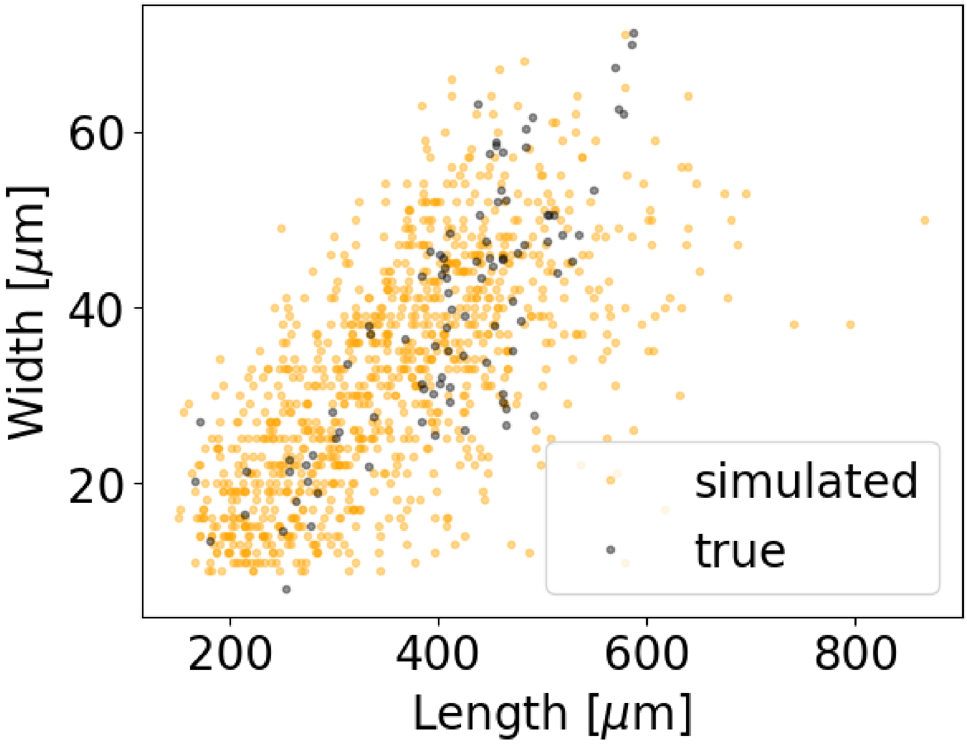


**Fig. S2** Distribution of lengths and widths of simulated and real worms.

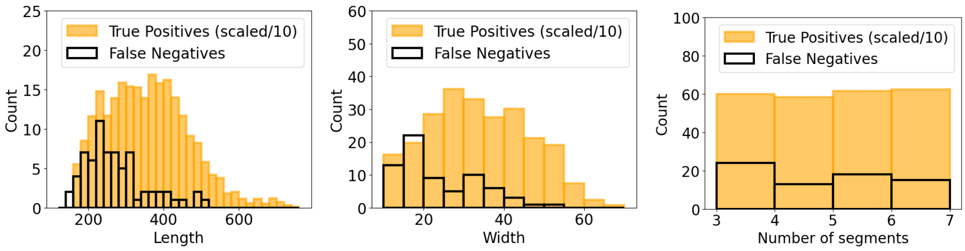


**Fig. S3** Histogram distributions of detections and non-detections as a function of various simulated nematode physical characteristics. The counts of detections have been reduced by a factor of 10 for visualization purposes.

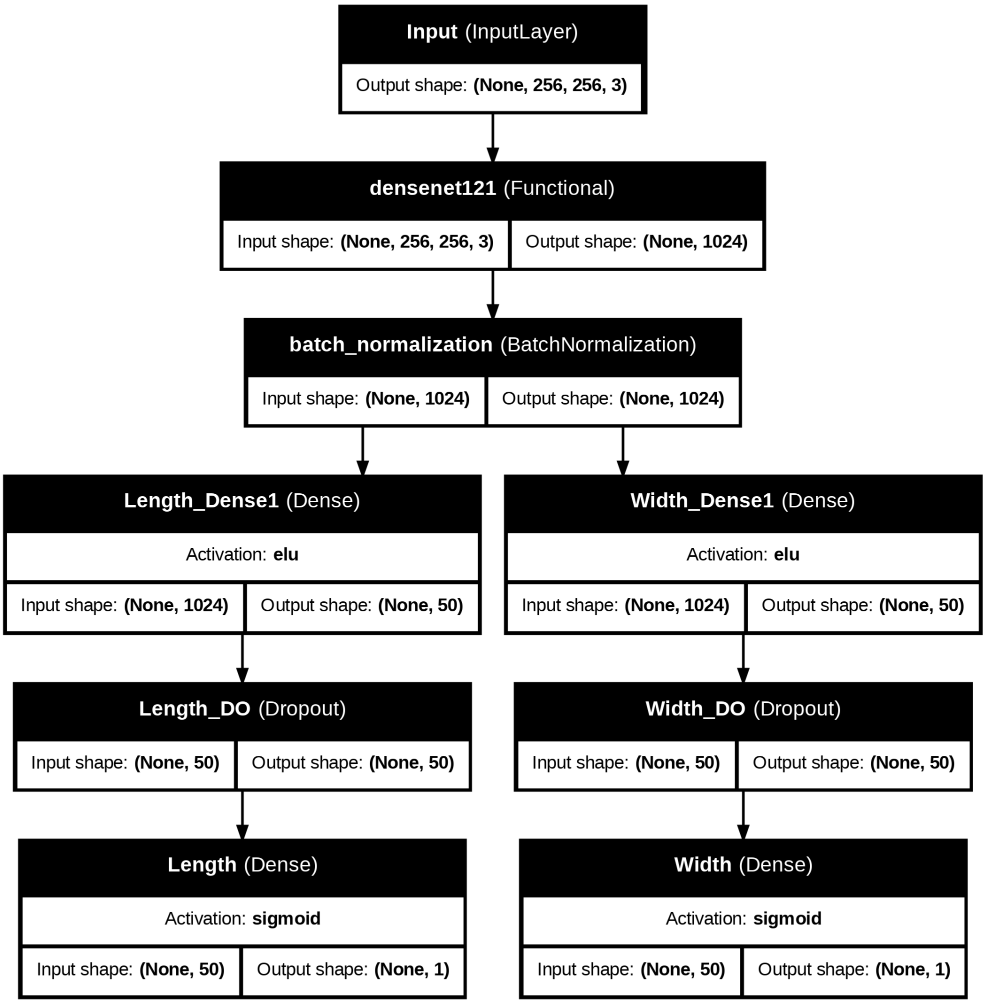


**Fig. S4** WormREG model architecture.
